# Re-examining the relationship between virus and microbial cell abundances in the global oceans

**DOI:** 10.1101/025544

**Authors:** Charles H. Wigington, Derek Sonderegger, Corina P.D. Brussaard, Alison Buchan, Jan F. Finke, Jed Fuhrman, Jay T. Lennon, Mathias Middelboe, Curtis A. Suttle, Charles Stock, William H. Wilson, K. Eric Wommack, Steven W. Wilhelm, Joshua S. Weitz

**Affiliations:** School of Biology, Georgia Institute of Technology, Atlanta, GA 30332; Department of Mathematics and Statistics, Northern Arizona University, Flagstaff, AZ; Department of Biological Oceanography, Royal Netherlands Institute for Sea Research (NIOZ), Texel, The Netherlands; Department of Aquatic Microbiology, Institute for Biodiversity and Ecosystem Dynamics (IBED), University of Amsterdam, The Netherlands; Department of Microbiology, University of Tennessee-Knoxville, Knoxville, TN; Department of Earth, Ocean and Atmospheric Sciences, University of British Columbia, Vancouver BC V6T 1Z4; Dept of Biological Sciences, University of Southern California, Los Angeles 90089; Department of Biology, Indiana University, Bloomington, IN 47405; Marine Biological Section, Department of Biology, University of Copenhagen, Helsingr, Denmark; Departments of Earth, Ocean and Atmospheric Sciences, Microbiology and Immunology, and Botany, University of British Columbia, Vancouver BC V6T 1Z4; Program in Integrated Microbial Diversity, Canadian Institute for Advanced Research, Toronto ON M5G 1Z8; Geophysical Fluid Dynamics Laboratory Princeton, NJ 08540-6649; Sir Alister Hardy Foundation for Ocean Science, The Laboratory, Citadel Hill, Plymouth, UK; Plant and Soil Sciences Delaware Biotechnology Institute Newark, DE; School of Physics, Georgia Institute of Technology, Atlanta, GA 30332

## Abstract

Marine viruses are critical drivers of ocean biogeochemistry and their abundances vary spatiotemporally in the global oceans, with upper estimates exceeding 10^8^ per ml. Over many years, a consensus has emerged that virus abundances are typically 10-fold higher than prokaryote abundances. The use of a fixed-ratio suggests that the relationship between virus and prokaryote abundances is both predictable and linear. However, the true explanatory power of a linear relationship and its robustness across diverse ocean environments is unclear. Here, we compile 5671 prokaryote and virus abundance estimates from 25 distinct marine surveys to characterize the relationship between virus and prokaryote abundances. We find that the median virus-to-prokaryote ratio (VPR) is 10:1 and 16:1 in the near-and sub-surface oceans, respectively. Nonetheless, we observe substantial variation in the VPR and find either no or limited explanatory power using fixed-ratio models. Instead, virus abundances are better described as nonlinear, power-law functions of prokaryote abundances - particularly when considering relationships within distinct marine surveys. Estimated power-laws have scaling exponents that are typically less than 1, signifying that the VPR decreases with prokaryote density, rather than remaining fixed. The emergence of power-law scaling presents a challenge for mechanistic models seeking to understand the ecological causes and consequences of marine virus-microbe interactions. Such power-law scaling also implies that efforts to average viral effects on microbial mortality and biogeochemical cycles using “representative” abundances or abundance-ratios need to be refined if they are to be utilized to make quantitative predictions at regional or global ocean scales.

---

Viruses of microbes have been linked to central processes across the global oceans, including biogeochemical cycling [9, 29, 39, 43, 47, 53] and the maintenance and generation of microbial diversity [3, 36, 39, 47, 52]. Virus propagation requires contacting and infecting cells. The per cell rate at which microbial cells – including bacteria, archaea, and microeukaryotes – are contacted by viruses is assumed to be proportional to the product of virus and microbial abundances [33]. If virus and microbe abundances were related in a predictable way it would be possible to infer the rate of contact, and potentially the relative importance of virus-induced cell lysis, from estimates of microbial abundance alone.

Virus ecology underwent a transformation in the late 1980s with the recognition that virus abundances, as estimated using culture-independent methods, were orders of magnitude higher than estimates via culture-based methods [4]. Soon thereafter, researchers began to report the “virus to bacterium ratio” (VBR) as a statistical proxy for the strength of the relationship between viruses and their potential hosts in both freshwater and marine systems [31]. This ratio is more appropriately termed the “virus-to-prokaryote ratio” (VPR) – a convention which we use here [1].

Observations accumulating over the past 25 years have observed wide variation in VPR, yet there is an emergent census that a suitable first-approximation is that VPR is 10 (see Table I). This ratio also reflects a consensus that typical prokaryote abundances are approximately 10^6^ per ml and typical virus abundances are approximately 10^7^ per ml [50, 57]. Yet, the use of a fixed ratio carries with it another assumption: that of linearity, i.e., if prokaryote abundance were to double, then viruses are expected to double as well. An alternative is that the relationship between virus and prokaryote abundance is better described in terms of a nonlinear relationship, e.g., a power-law.

**TABLE I:**
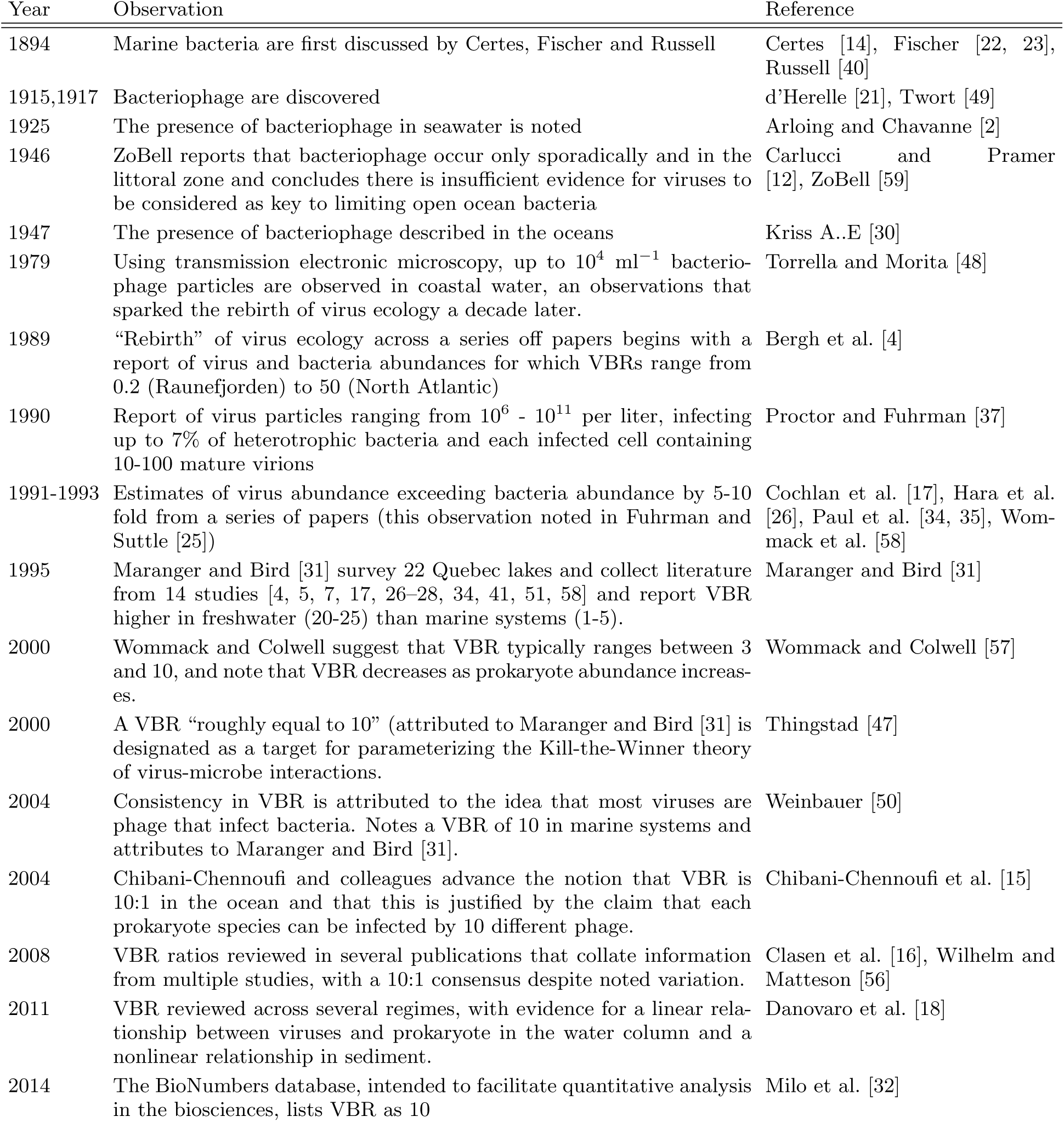
Origins and emerging consensus of the 10:1 ratio of virus abundance to bacteria abundance in aquatic systems - from freshwater lakes to the global oceans. We use the convention VPR in this mansucript rather than VBR.

| Year | Observation | Reference |
| --- | --- | --- |
| 1894 | Marine bacteria are first discussed by Certes, Fischer and Russell | Certes [14], Fischer [22, 23], Russell [40] |
| 1915,1917 | Bacteriophage are discovered | d’Herelle [21], Twort [49] |
| 1925 | The presence of bacteriophage in seawater is noted | Arloing and Chavanne [2] |
| 1946 | ZoBell reports that bacteriophage occur only sporadically and in the littoral zone and concludes there is insufficient evidence for viruses to be considered as key to limiting open ocean bacteria | Carlucci and Pramer [12], ZoBell [59] |
| 1947 | The presence of bacteriophage described in the oceans | Kriss A..E [30] |
| 1979 | Using transmission electronic microscopy, up to $10^4$ ml<sup>-1</sup> bacteriophage particles are observed in coastal water, an observations that sparked the rebirth of virus ecology a decade later. | Torrella and Morita [48] |
| 1989 | “Rebirth” of virus ecology across a series off papers begins with a report of virus and bacteria abundances for which VBRs range from 0.2 (Raunefjorden) to 50 (North Atlantic) | Bergh et al. [4] |
| 1990 | Report of virus particles ranging from $10^6$ - $10^{11}$ per liter, infecting up to 7% of heterotrophic bacteria and each infected cell containing 10-100 mature virions | Proctor and Fuhrman [37] |
| 1991-1993 | Estimates of virus abundance exceeding bacteria abundance by 5-10 fold from a series of papers (this observation noted in Fuhrman and Suttle [25]) | Cochlan et al. [17], Hara et al. [26], Paul et al. [34, 35], Wommack et al. [58] |
| 1995 | Maranger and Bird [31] survey 22 Quebec lakes and collect literature from 14 studies [4, 5, 7, 17, 26-28, 34, 41, 51, 58] and report VBR higher in freshwater (20-25) than marine systems (1-5). | Maranger and Bird [31] |
| 2000 | Wommack and Colwell suggest that VBR typically ranges between 3 and 10, and note that VBR decreases as prokaryote abundance increases. | Wommack and Colwell [57] |
| 2000 | A VBR “roughly equal to 10” (attributed to Maranger and Bird [31] is designated as a target for parameterizing the Kill-the-Winner theory of virus-microbe interactions. | Thingstad [47] |
| 2004 | Consistency in VBR is attributed to the idea that most viruses are phage that infect bacteria. Notes a VBR of 10 in marine systems and attributes to Maranger and Bird [31]. | Weinbauer [50] |
| 2004 | Chibani-Chennoufi and colleagues advance the notion that VBR is 10:1 in the ocean and that this is justified by the claim that each prokaryote species can be infected by 10 different phage. | Chibani-Chennoufi et al. [15] |
| 2008 | VBR ratios reviewed in several publications that collate information from multiple studies, with a 10:1 consensus despite noted variation. | Clasen et al. [16], Wilhelm and Matteson [56] |
| 2011 | VBR reviewed across several regimes, with evidence for a linear relationship between viruses and prokaryote in the water column and a nonlinear relationship in sediment. | Danovaro et al. [18] |
| 2014 | The BioNumbers database, intended to facilitate quantitative analysis in the biosciences, lists VBR as 10 | Milo et al. [32] |

In practice, efforts to predict the regional or global-scale effects of viruses on marine microbial mortality, turnover and even biogeochemical cycles, depend critically on the predictability of the relative density of viruses and microbial cells. Here, we directly query the nature of the relationship between viruses and prokaryotes via a large-scale compilation and re-analysis of abundance data across marine environments.

## I. RESULTS

### A. VPR exhibits substantial variation in the global oceans

In the compiled marine survey data (see Figure 1, Table S1 and the Materials and Methods), 95% of prokaryote abundances range from 5.0 ×10^3^ to 4.1 × 10^6^ per ml and 95% of virus abundances range from roughly 3.7 × 10^5^ to 6.4 × 10^7^ per ml (Figure 2A). Both prokaryote and virus concentrations generally decrease with depth as reported previously (e.g., see [18]). The median VPR for the near-surface samples (*≤* 100m) is 10.5 and the median VPR for the sub-surface samples (*>* 100m) is 16.0. In that sense, the consensus 10:1 ratio does accurately represent the median VPR for the surface data. We also observe substantial variation in VPR, as has been noted in prior surveys and reviews (see Table 1). Figure 2B shows that 95% of the variation in VPR in the near-surface ocean lies between 1.4 and 160 and between 3.9 and 74 in the sub-surface ocean. For the near-surface ocean, 44% of the VPR values are between 5 and 15, 16% are less than 5 and 40% exceed 15. This wide distribution, both near-and sub-surface demonstrates potential limitations in utilizing the 10:1 VPR, or any fixed ratio, as the basis for a *predictive* model of virus abundance derived from estimates of prokaryote abundance.

**FIG. 1:**
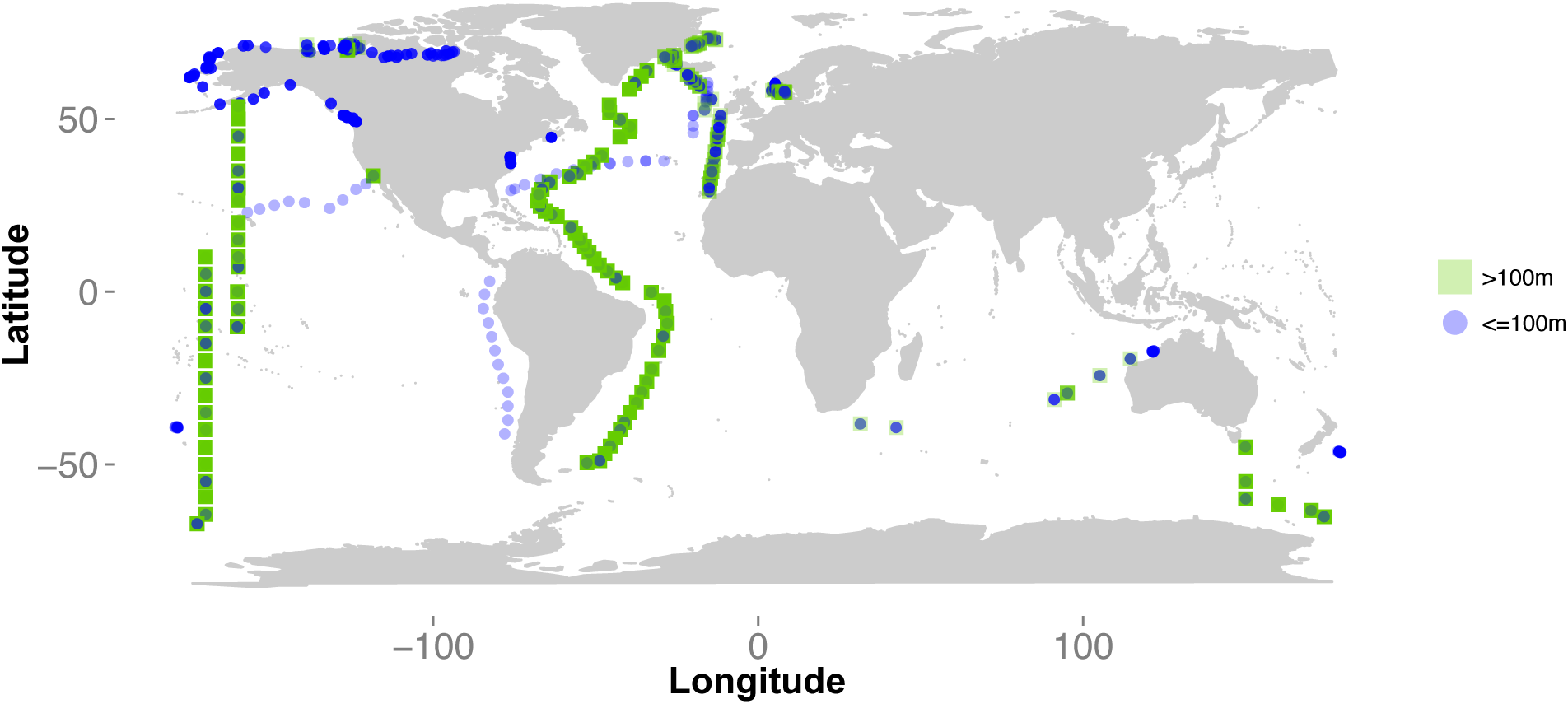
Global distribution of sample sites. Each point denotes a location from which one or more samples were taken. Samples range from the surface to up to 5,500 meters below sea level, with 2,921 taken near the surface (*≤* 100m), noted as squares, and 2,750 taken below the surface (*>*100m), noted as circles. The number of points for each study – the “Frequency” – is found in Table S2. Full access to data is provided as Supplementary File 1.

**FIG. 2:**
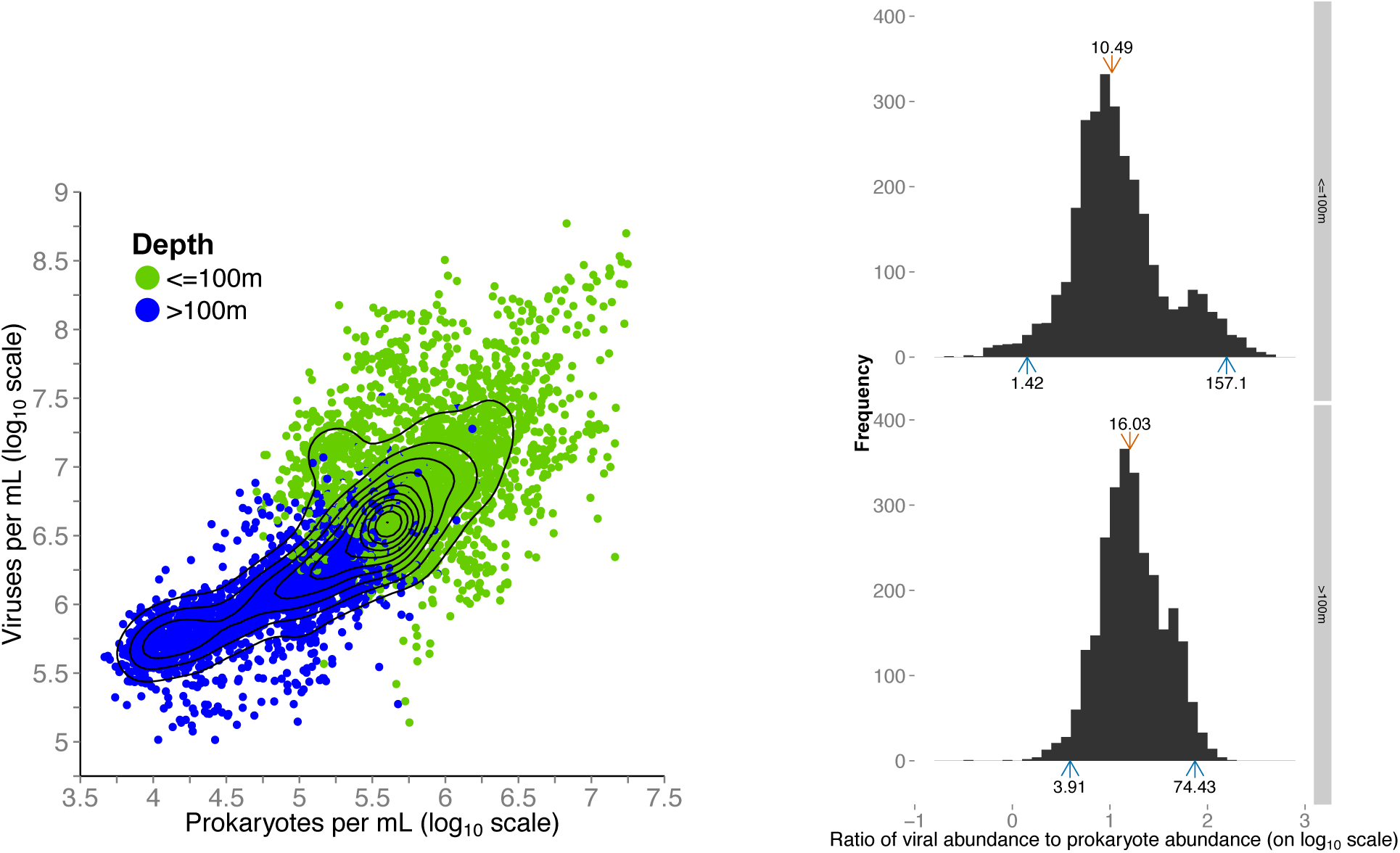
Variation in virus and prokaryote abundances and the VPR. (A) Prokaryote abundance vs. virus abundance colored by depth. Each point represents a biological sample. (B) Histogram of the logarithm of VPR. The top and bottom panels correspond to near- and sub-surface water column samples, respectively. The red arrow denotes the median value and the blue arrows denote the central 95% range of values - where the numbers associated with each arrow denote the non-transformed value of VPR.

### B. Virus abundance does not vary linearly with prokaryote abundance

Figure 3 shows two alternative, predictive models of the relationship between logarithmically scaled virus and prokaryote abundances for water column samples. The models correspond to a fixed-ratio model and a power-law model. To clarify the interpretation of fitting in log-log space consider a fixed-ratio model with a 12:1 ratio between virus and prokaryote abundance, *V* = 12 *≤ B*. Then, in log-log space the relationship is

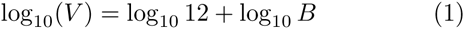

**FIG. 3:**
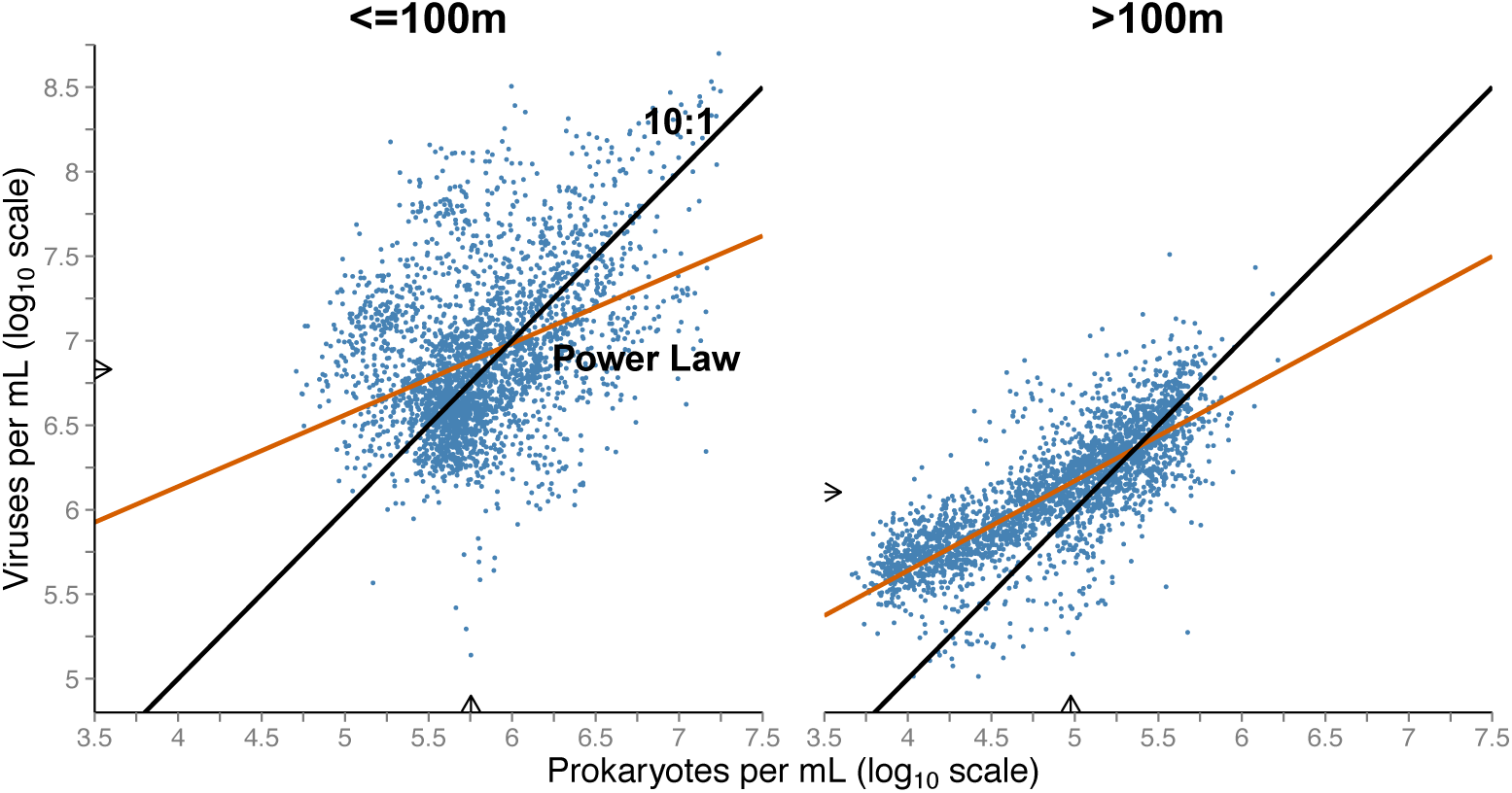
Virus abundance is poorly fit by a model of 10-fold increase relative to prokaryote abundance. (Left) Surface ocean – the red line denotes the best fit power-law with an exponent of 0.42 while the black line denotes the 10:1 curve. The best-fit power law explains 15% of the variation and the 10:1 line explains -16% of the variation. See text for interpretation of negative *R*^2^ and the importance of outliers in these fits. (Right) Deeper water column – the red line denotes the best fit power-law with an exponent of 0.53 while the black line denotes the 10:1 curve. The best-fit power law explains 64% of the variation and the 10:1 line explains -26% of the variation In both cases the arrows on the axes denote the median of the respective abundances.

which we interpret as a line with y-intercept of log_10_ 12 = 1.08 and a slope (change in *log*_10_*V* for a 1-unit change in log_10_ *B*) of 1. By the same logic, any fixed-ratio model will result in a line with slope 1 in the log-log plot and the y-intercept will vary logarithmically with VPR. The alternative predictive model is that of a power-law: *V* = *cB*^*α*1^. In log-log space, the relationship is:

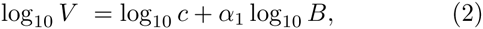

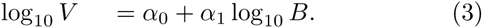

The slope, *α*_1_, of a fitted line on log-transformed data denotes the *power-law exponent* that best describes the relationship between the variables. The intercept, *α*_0_, of a fitted line on log-transformed data denotes the logarithmically transformed pre factor.

The 10:1 line has a residual squared error of -16% and -25% in the surface and deep samples, respectively (Table 2). In both cases, this result means that a 10:1 line explains *less* of the variation in virus abundance compared to a model in which virus abundance is predicted by its mean value across the data. In order to evaluate the generality of this result, we considered an ensemble of fixed-ratio models each with a different VPR. In the near-surface samples, we find that all fixed-ratio models explain less of the variation (i.,e., have *negative* values of *R*^2^) than does a “model” in which virus abundance is predicted to be the global mean in the dataset (Figure S1). This reflects the failure of constant ratio (i.e., linear) models to capture the cluster of high VPRs at low prokaryote density apparent in the density contours of Figure 2A and the shoulder of elevated high VPR frequency in Figure 2B. The largest contributor to this cluster of points is the Arctic SBI study (see Figure S1). Whereas, in the sub-surface samples, fixed-ratio models in which VPR varies between 12 and 22 do have positive explanatory power, but all perform worse than does the power-law model (Figure S1). In contrast, the best fitting power-law model explains 15% and 64% of the variation in the data, for near-and sub-surface samples respectively (Table 2). The best-fit power-law scaling exponent is 0.42 with 95% confidence intervals (CIs) of (0.39, 0.46) for near-surface samples and 0.53 with 95% CIs of (0.52, 0.55) for sub-surface samples. In other words, doubling prokaryote abundance along either regression line is not expected to lead to a doubling in virus abundance, but rather a 2^0.42^ = 1.33 and 2^0.53^ = 1.44 fold increase, respectively. The power-law model is an improvement over the fixed ratio model in both cases, even when accounting for the increase in parameters (Table 2). In the near-surface, refitting surface data without outliers improves explanatory power to approximately *R*^2^ = 0.3 in contrast to an *R*^2^ = 0.65 for the sub-surface (see Methods and Figure S2). In summary, the predictive value of a power-law model is much stronger in the sub-surface than in the near-surface, where confidence in the interpretation of power-law exponents is limited.

**TABLE II:** Information theoretic comparison of alternative models of the relationship between virus and prokaryote abundance. The values of the Aikake Information Criteria (AIC) are defined in the Materials and Materials and Methods. The value of *R*^2^ for each model denotes the relative amount of variance explained. Negative values of *R*^2^ mean that a model explains less variance than does the overall mean.

| Model | ≤ 100m |  | >100m |  |
| --- | --- | --- | --- | --- |
| | $R^2$ | AIC | $R^2$ | AIC |
| 10:1 | -0.16 | -15305.83 | -0.25 | -14492.09 |
| Power Law | 0.15 | -16301.81 | 0.64 | -18313.82 |
| Constrained Power Law | 0.39 | -17292.11 | 0.66 | -18513.48 |
| Power Law by Study | 0.79 | -20293.10 | 0.72 | -18972.81 |

### C. Study-to-study measurement variation is unlikely to explain the intrinsic variability of virus abundances in the surface ocean

Next, we explored the possibility that variation in methodologies affected the baseline offset of virus abundance measurements and thereby decreased the explanatory power of predicting virus abundances based on prokaryote abundances. That is, if *V \** is the true and unknown abundance of viruses, then it is possible that two studies would estimate 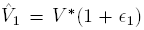 and 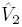 = *V*\*(1 + *ε*_2_) where |***ϵ***_1_| and |***ϵ***_2_| denote the relative magnitude of study-specific shifts. We constrain the relative variation in measurement, such that the measurement uncertainty is 50% or less (see Materials and Methods). The constrained regression model improves the explanatory power of the model (see Table 2), but in doing so, the model forces 18 of the 25 studies to the maximum level of measurement variation permitted (Figure S3). We do not expect that differences in measurement protocols to explain nearly 2 orders of magnitude variation in estimating virus abundance given the same true virus abundance in a sample. Note that when sub-surface samples were analyzed through the constrained power-law model, there was only a marginal increase of 2% in *R*^2^ and, moreover, 9 of the 12 studies were fit given the maximum level of measurement variation permitted (Figure S3)[h!]. The constrained intercept model results suggest that the observed variation in virus abundance in the surface oceans is not well explained strictly by variation in measurement protocol between studies.

### D. VPR decreases with increasing prokaryote abundance – a hallmark of power-law relationships

We next evaluate an ensemble of power-law models: *V*_*i*_ = *c*_*i*_*N*^α_*i*_^ where the index *i* denotes the use of distinct intercepts and power-law exponents for each survey. The interpretation of this model is that the non-linear nature of the virus to prokaryote relationship may differ in distinct oceanic realms or due to underlying differences in sites or systems, rather than due to measurement differences. Figure 4 shows the results of fitting using the study-specific power-law model in the surface ocean samples. Study-specific power-law fits are significant in 18 of 25 cases in the surface ocean. The median power-law exponent for studies in the surface ocean is 0.50. Furthermore, of those significant power-law fits, the 95% distribution of the power-law exponent excludes a slope of one and is entirely less than one in 11 of 18 cases (see Figure 5). This model in which the power-law exponent varies with study is a significant improvement in terms of *R*^2^ (Table 2). For sub-surface samples, study-specific power-law fits are significant in 10 of 12 cases in the sub-surface (Figure S4). The median power-law exponent for studies in the sub-surface is 0.67. Of those significant power-law fits, the 95% distribution of the power-law exponent is entirely less than one in 6 of 10 cases (see Figure S5). A power-law exponent of less than one means that virus abundance increases less than proportionately given increases in prokaryote abundance. This study-specific analysis extends the findings that nonlinear, rather than linear, models are more suitable to describe the relationship between virus and prokaryote abundances. We find that the dominant trend in both near-surface and sub-surface samples is that VPR decreases as prokaryote abundance increases. The increased explanatory power by study is stronger for near-surface than for sub-surface samples. This increase in *R*^2^ comes with a caveat: study-specific models do not enable *a priori* predictions of virus abundance given a new environment or sample site, without further effort to disentangle biotic and abiotic factors underlying the different scaling relationships.

**FIG. 4:**
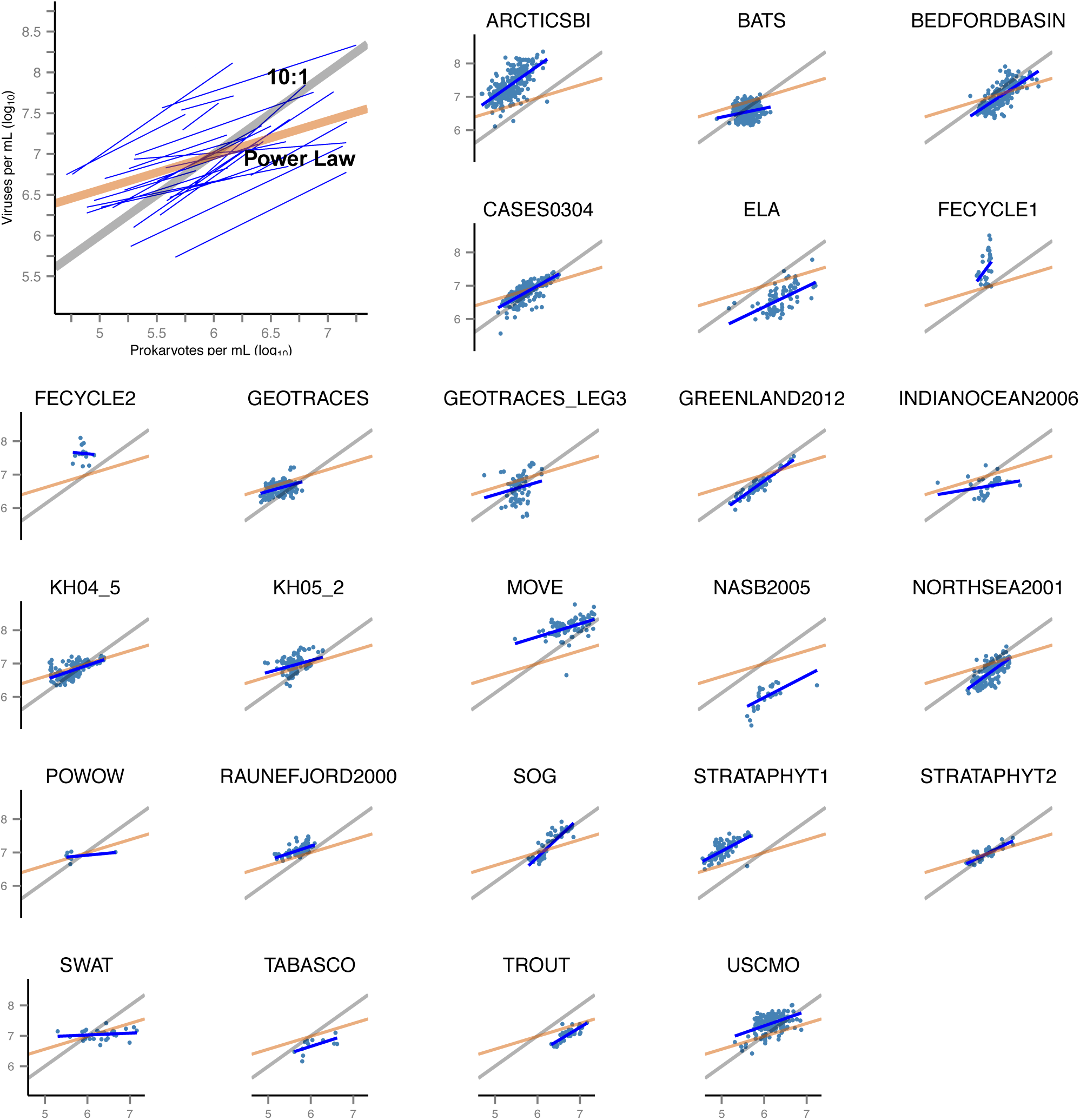
Virus-prokaryote relationships given the variable slope and intercept mixed-effects model. (Upper-left) Best-fit power-law for each study (blue lines) plotted along with the best-fit power-law of the entire dataset (red line) and the 10:1 line (grey line). (Individual panels) Best-fit power-law model (blue line) on log-transformed data (blue points) for each study, with the power-law model regression (red) and 10:1 line (black) as reference. The power-law exponents and associated confidence intervals are shown in Figure 5,

**FIG. 5:**
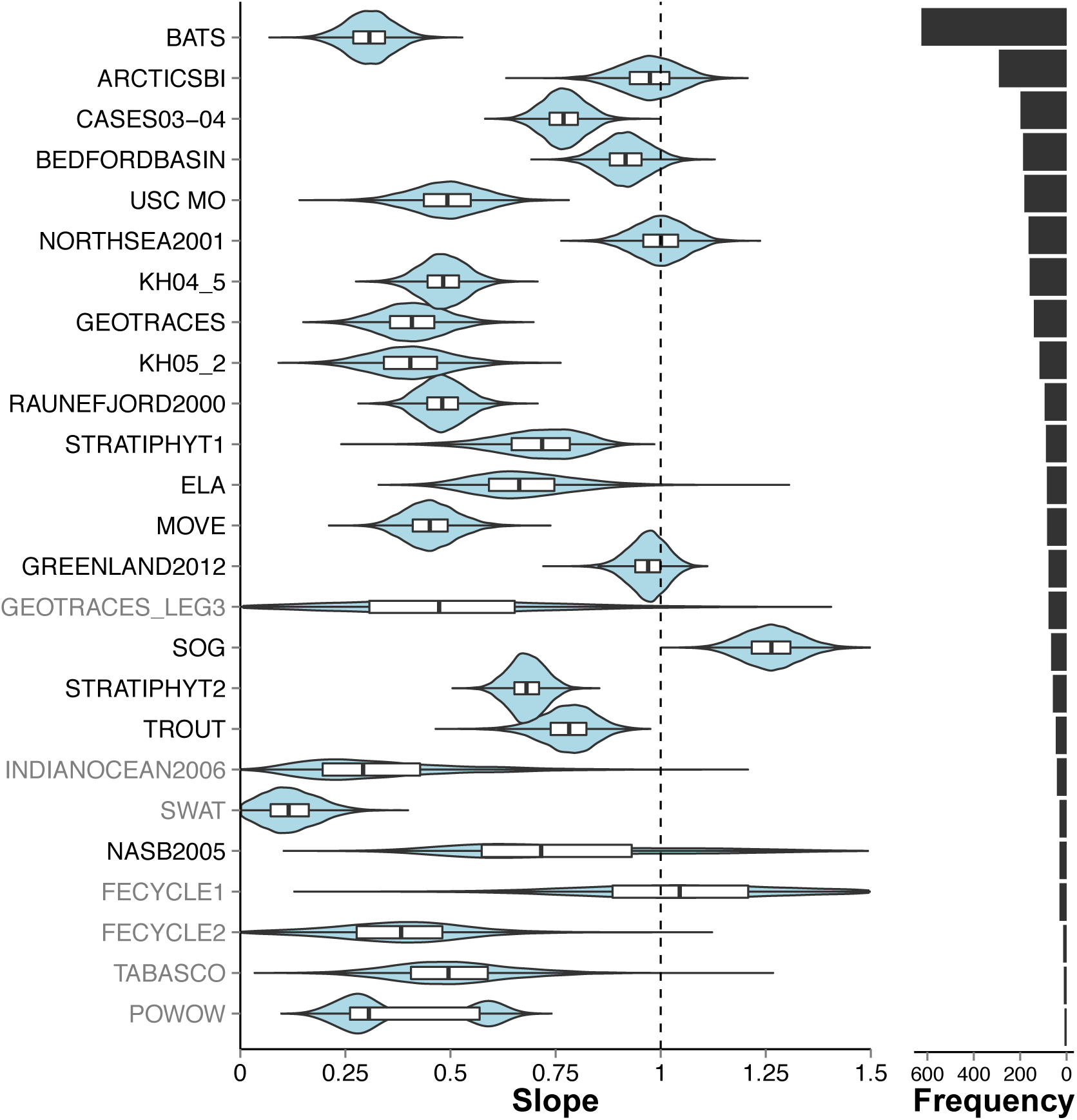
Study-specific 95% confidence intervals of power-law exponents for relationships between virus and prokaryote abundance in the surface. The confidence intervals are plotting using “violin” plots including the median (center black line), 75% distribution (white bars) and 95% distribution (black line), with the distribution overlaid (blue shaded area). The number of points included as part of each study is displayed on the right-most bar plots. Study labels in black indicate those studies for which the regression fit had a p-value less than 0.002=0.05/25 (accounting for a multiple comparison correction given the analysis of 25 studies). Study labels in gray indicate a p-value above this threshold.

## II. DISCUSSION

Viruses are increasingly considered as part of efforts to understand the factors controlling marine microbial mortality, productivity, and biogeochemical cycles [8, 9, 19, 39, 45, 55]. Quantitative estimates of these viral-induced effects can be measured directly, but are often inferred indirectly, using the relative abundance of viruses to prokaryotes. To do so, there is a consensus that assuming the virus-to-prokaryote ratio is 10 in the global oceans - despite observed variation - is a reasonable starting point. Here, we have re-analyzed the relationship of virus to prokaryote abundance in 25 marine survey datasets. We find that 95% of the variation in VPR ranges from 1.4 to 160 in the near-surface ocean and from 3.9 to 74 in the sub-surface. Although the 10:1 ratio accurately describes the median of the VPR in the surface ocean, the broad distribution of VPR implies that prokaryote abundance is a poor quantitative predictor of virus abundance. Moreover, increases in prokaryote abundance do not lead to proportionate increases in virus abundance. Instead, we propose that the virus to prokaryote abundance relationship is nonlinear, and that the degree of nonlinearity – as quantified via a power-law exponent – is typically less than 1. This sublinear relationship can be interpreted to mean that VPR decreases as an increasing function of prokaryote abundance, and generalizes earlier observations [57].

Power-law relationships between virus and prokaryote abundance emerge from complex feedbacks involving both exogeneous and endogenous factors. The question of exogenous factors could be addressed, in part, by examining environmental covariates at survey sites. For example, if prokaryote and virus abundances varied systematically with another environmental co-factor during a transect, then this would potentially influence the inferred relationship between virus and prokaryote abundances. In that same way, variation in environmental correlates, including temperature and incident ration, may directly modify virus life history traits [20, 44]. Likewise, some of the marine survey datasets examined here constitute repeated measurements at the same location (e.g., at the Bermuda Atlantic Time-series Study (BATS)). Time-varying environmental factors could influence the relative abundance of microbes and viruses. It is also interesting to note that viruses-induced mortality is considered to be more important at eutrophic sites [57], where microbial abundance is higher - yet the observed decline in VPR with prokaryote abundance would suggest the opposite.

It could also be the case that variation in endogenous factors, e.g., the life history traits of viruses and microbes, including variability in the structure of cross-infection [54], could drive changes in emergent relationships between virus and prokaryote abundances. Virus-microbe interactions can lead to intrinsic oscillatory dynamics. Indeed, previous observations of a declining relationship between VPR and prokaryote abundance have been attributed to changing ratios across phytoplankton bloom events, including possible virus-induced termination of blooms [57]. Similar arguments were proposed in the analysis of tidal sediments [13]. Alternatively, observations of declining VPR with prokaryote density have been attributed to variation in underlying diversity [6]. Whatsoever the mechanism(s), it is striking that virus abundances in some surveys can be strongly predicted via alternative power-law functions of prokaryote abundances. Mechanistic models are needed to further elucidate these emergent macroecological patterns and relationships.

The present analysis separated the abundance data first according to depth and then according to survey as a means to identify different relationships between virus and prokaryote abundances in the global oceans. The predictive value of total prokaryote abundance is strong when considering sub-surface samples. In contrast, prokaryote abundance is not a strong predictor of virus abundance in the near-surface samples, when utilizing linear or nonlinear models. The predictive power of nonlinear models improved substantially in the near-surface when evaluating each marine survey separately. The minimal predictive value of prokaryoate abundances in the near-surface when aggregating across all surveys is problematic given that virus-microbe interactions have significant roles in driving prokaryote mortality and ecosystem functioning [9, 45, 53]. Indeed the aggregation of abundance measurements in terms of total prokaryote abundances may represent part of the problem. At a given site and time of sampling, each microbial cell in the community is potentially targeted by a subset of the total viral pool. In moving forward, understanding variation in virus abundance and its relationship to prokaryote abundance requires a critical examination of correlations at functionally relevant temporal and spatial scales, i.e., at the scale of interacting pairs of viruses and microbes. We encourage the research community to prioritize examination of these scales of interaction as part of efforts to understand mechanisms underlying nonlinear virus-microbe abundance relationships in the global oceans.

## III. MATERIALS AND METHODS

### A. Data source

Marine virus abundance data was aggregated from 25 studies (Table S1). A total of 5671 data points were aggregated. The data collection dates range from 1996 to 2012. Data primarily comes from coastal waters in the northern hemisphere and were collected predominately during the summer months, with the notable exceptions of long-term coastal monthly monitoring sites, i.e., the studies USC MO, BATS, and MOVE.

### B. Data processing

Analyses of the data were performed using R version 3.1.1. Scripts and original data are provided at https://github.com/cwigington3/VIRBAC analysis.

### C. Power-law model

A power-law regression model used the *log*_10_ of the predictor variable, prokaryote abundance per mL *N*, and the *log*_10_ of the outcome variable, virus abundance per mL *V*. The power-law regression was calculated using the equation log_10_ *V* = *α*_0_ + *α*_1_ log_10_ *N*. The *α*_0_ and *α*_1_ parameters were fit via OLS regression to minimize the sum of square error.

### D. Constrained variable-intercept model

The constrained model is a “mixed-effects” regression model using the same predictor and outcome variables, *log*_10_ of prokaryote abundance per mL and the *log*_10_ virus abundance per mL, respectively. This model includes study-specific intercepts which were constrained such that the value for any of the intercepts were restricted to one standard error above or below the intercept value taken from the power-law model. The standard error value for this model came from the power-law model. The equation for this model is 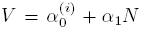, where 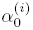 is the study-specific intercept and and *α*_1_ is the slope common to all studies, *N* is the predictor variable, and *V* is the outcome variable.

### E. Variable slope and variable intercept model

A power-law model where the exponent and intercept varied with each study was evaluated using the same predictor variable, log_10_ prokaryote abundance per mL, and the same outcome variable, log_10_ virus abundance per mL. In this model, there was a study-specific *α*_0_ and *α*_1_ and a OLS regression calculated using the equation V = 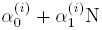.

### F. Bootstrapping model confidence intervals

Bootstrap analyses of the power-law model and mixed effects models were conducted to derive 95% confidence intervals surrounding the parameters estimated by the models. For all models the original dataset was sampled with replacement, by study, to arrive at a bootstramp sample dataset, this process was repeated 10,000 times. Distributions for all parameters were generated and the 2.5%, 50%, and 97.5% points were identified from among the 10,000 parameter estimates.

### G. Outlier identification

Outliers in the data were identified by calculating the top and bottom 2% of estimated VPR amongst the entire 5,671 samples. The outliers corresponded to ratios below 1.81 and above 128. Those samples with virus to prokaryote ratios which fell outside of these bounds were considered outliers. There were 218 outlier samples taken at depths *≤* 100m and 10 outlier samples taken at depths *>*100m.

This work was supported by NSF grants OCE-1233760 (JSW) and OCE-1061352 (AB and SWW), a Career Award at the Scientific Interface from the Burroughs Wellcome Fund (JSW) and a Simons Foundation SCOPE grant (JSW). This work arose from discussions in the Ocean Viral Dynamics working group at the National Institute for Mathematical and Biological Synthesis, an Institute sponsored by the National Science Foundation, the U.S. Department of Homeland Security, and the Department of Agriculture through NSF Award EF-0832858, with additional support from The University of Tennessee, Knoxville.

---

**TABLE S1:** Virus and microbial abundance data from 25 different marine virus abundance studies from 11 different lab groups. A total of 5671 data points were aggregated. The data collection dates range from 2000 to 2011. Due to sampling convenience, data primarily comes from coastal waters in the northern hemisphere and were collected predominately during the summer months, with the notable exceptions of long-term coastal monthly monitoring sites (USC MO,BATS,Cheasapeake Bay).

| Study Name | Lab | Study Type | Location | Regime | Citation |
| --- | --- | --- | --- | --- | --- |
| NORTHSEA2001 | Bratback | Spatial | North Sea | Coastal | Bratback (unpublished) |
| RAUNEFJORD2000 | Bratback | Temporal | North Sea | Coastal | Bratback (unpublished) |
| BATS | Breitbart | Temporal | Sargasso Sea | nonCoastal | Parson et al. 2011 |
| STRATIPHYT1 | Brussaard | Spatial | N-Atlantic Transect | nonCoastal | Mojica et al., 2015 |
| STRATIPHYT2 | Brussaard | Spatial | N-Atlantic Transect | nonCoastal | Brussaard (unpublished) |
| USC MO | Fuhrman | Temporal | Santa Barbara Channel | nonCoastal | Fuhrman et al. 2006 |
| GEOTRACES | Herndl | Spatial | Atlantic Transect | nonCoastal | De Corte et al. 2012 |
| GEOTRACES.LEG3 | Herndl | Spatial | Atlantic Transect | nonCoastal | Herndl (unpublished) |
| BEDFORDBASIN | Li | Temporal | North Atlantic Ocean | Coastal | Li and Dickie 2001 |
| GREENLAND 2012 | Middleboe | Spatial | Greenland Sea | nonCoastal | Middleboe (unpublished) |
| INDIANOCEAN2006 | Middleboe | Spatial | Indian Ocean | nonCoastal | Middleboe (unpublished) |
| KH04-5 | Nagata | Spatial | Southern Pacific Ocean | nonCoastal | Yang et al. 2014 |
| KH05-2 | Nagata | Spatial | Northern Pacific Ocean | nonCoastal | Yang et al. 2014 |
| CASES03-04 | Suttle | Spatial | Arctic Ocean | nonCoastal | Payet and Suttle, 2013 |
| SOG/ELA/TROUT/SWAT | Suttle | Temporal | Pacific Ocean - Strait of Georgia | Coastal | Clasen et al., 2008 |
| ARCTICSBI | Wilhelm | Spatial | Gulf of Alaska | Coastal | Balsom, MS thesis 2003 |
| FECYCLE2 | Wilhelm | Spatial | South Pacific Ocean | nonCoastal | Matteson et al. 2012 |
| FECYCLE1 | Wilhelm | Spatial | South Pacific Ocean | nonCoastal | Stzepek et al. 2005 |
| MOVE | Wommack | Temporal | Atlantic - Chesapeake | Coastal | Wang et al. 2011 |
| NASB2005 | Wilhelm | Spatial | North Atlantic Ocean | nonCoastal | Rowe et al. 2008 |
| POWOW | Wilhelm | Spatial | Pacific Ocean | nonCoastal | Wilhelm (unpublished) |
| TABASCO | Wilhelm | Spatial | South Pacific Ocean | nonCoastal | Wilhelm et al. 2003 |

**TABLE S2:** Number of data points per study.

| Study | $\leq 100\text{m}$ | $> 100\text{m}$ | Total |
| --- | --- | --- | --- |
| ARCTICSBI | 292 | 0 | 292 |
| BATS | 626 | 756 | 1382 |
| BEDFORDBASIN | 188 | 0 | 188 |
| CASES03-04 | 199 | 46 | 245 |
| ELA | 85 | 0 | 85 |
| FECYCLE1 | 31 | 0 | 31 |
| FECYCLE2 | 15 | 0 | 15 |
| GEOTRACES | 141 | 631 | 772 |
| GEOTRACES_LEG3 | 78 | 351 | 429 |
| GREENLAND2012 | 78 | 46 | 124 |
| INDIANOCEAN2006 | 42 | 10 | 52 |
| KH04.5 | 159 | 383 | 542 |
| KH05.2 | 117 | 238 | 355 |
| MOVE | 84 | 0 | 84 |
| NASB2005 | 31 | 0 | 31 |
| NORTHSEA2001 | 164 | 27 | 191 |
| POWOW | 9 | 0 | 9 |
| RAUNEFJORD2000 | 95 | 0 | 95 |
| SOG | 67 | 0 | 67 |
| STRATIPHYT1 | 89 | 24 | 113 |
| STRATIPHYT2 | 59 | 34 | 93 |
| SWAT | 31 | 0 | 31 |
| TABASCO | 12 | 0 | 12 |
| TROUT | 47 | 0 | 47 |
| USC MO | 182 | 204 | 386 |
| Total | 2921 | 2750 | 5671 |

**TABLE S3:** Variation in the estimate of the intercept, 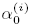, for each study and associated standard error for the constrained power-law model as applied to surface ocean data. The common intercept in this model is *α_0_* = 4.44 and the common slope is 0.42. The group column denotes whether the study-specific intercept exceeds that of the common intercept (denoted as group A) or is below that of the common intercept (denoted as group B). The table is sorted according to the lab-specific intercept estimates.

| Study | Intercept | Std. Error | Group |
| --- | --- | --- | --- |
| ARCTICSBI | 4.552594 | 0.1580157 | A |
| FECYCLE1 | 4.552594 | 0.1837236 | A |
| FECYCLE2 | 4.552594 | 0.1961546 | A |
| MOVE | 4.552594 | 0.1982541 | A |
| Raunefjord | 4.552594 | 0.1716625 | A |
| StratiphytI | 4.552594 | 0.1543207 | A |
| USCMO | 4.552594 | 0.1806831 | A |
| KH05_2 | 4.513041 | 0.1683358 | A |
| SOG | 4.480697 | 0.1869220 | A |
| POWOW | 4.408948 | 0.2060002 | B |
| StratiphytII | 4.389658 | 0.1765001 | B |
| KHO4 | 4.339907 | 0.1688285 | B |
| CASES0304 | 4.339902 | 0.1669256 | B |
| BEDFORDBASIN | 4.336256 | 0.1825271 | B |
| BATS | 4.332784 | 0.1647363 | B |
| ELA | 4.332784 | 0.1885707 | B |
| GEOTRACES | 4.332784 | 0.1580225 | B |
| GEOTRACES_LEG3 | 4.332784 | 0.1679548 | B |
| GREENLAND2012 | 4.332784 | 0.1759093 | B |
| INDIANOCEAN2006 | 4.332784 | 0.1820002 | B |
| NASB2005 | 4.332784 | 0.1876671 | B |
| NORTHSEA2001 | 4.332784 | 0.1770218 | B |
| SWAT | 4.332784 | 0.1945882 | B |
| TABASCO | 4.332784 | 0.2047635 | B |
| TROUT | 4.332784 | 0.2019569 | B |

**TABLE S4:** Explanatory power and significance of power-law fits for the model in which the power-law exponent is allowed to vary between studies. Empty cells in a row denote the absence of samples collected at depths *>* 100 m for the study denoted in the left-most column.

| Study | $\leq 100$ m | | $> 100$ m | |
| --- | --- | --- | --- | --- |
| | $R^2$ | $p$-value | $R^2$ | $p$-value |
| ARCTICSBI | 0.441 | <1e-05 |  |  |
| BATS | 0.045 | <1e-05 | 0.504 | <1e-05 |
| BEDFORDBASIN | 0.537 | <1e-05 |  |  |
| CASES03-04 | 0.541 | <1e-055 | 0.072 | 0.0718 |
| ELA | 0.343 | <1e-05 |  |  |
| FECYCLE | 0.146 | 0.0341 |  |  |
| FECYCLE2 | 0.004 | 0.813 |  |  |
| GEOTRACES | 0.163 | <1e-05 | 0.706 | <1e-05 |
| GEOTRACES_LEG3 | 0.043 | 0.0695 | 0.396 | <1e-05 |
| GREENLAND2012 | 0.868 | <1e-05 | 0.333 | 2.7e-05 |
| INDIANOCEAN2006 | 0.068 | 0.0955 | 0.288 | 0.11 |
| KH04.5 | 0.325 | <1e-05 | 0.703 | <1e-05 |
| KH05.2 | 0.122 | 0.000112 | 0.836 | <1e-05 |
| MOVE | 0.24 | <1e-05 |  |  |
| NASB2005 | 0.382 | 0.00021 |  |  |
| NORTHSEA2001 | 0.542 | <1e-05 | 0.51 | 2.85e-05 |
| POWOW | 0.136 | 0.329 |  |  |
| RAUNEFJORD2000 | 0.349 | <1e-05 |  |  |
| SOG | 0.788 | <1e-05 |  |  |
| STRATIPHYT1 | 0.448 | <1e-05 | 0.471 | 0.000214 |
| STRATIPHYT2 | 0.768 | <1e-05 | 0.731 | <1e-05 |
| SWAT | 0.026 | 0.389 |  |  |
| TABASCO | 0.371 | 0.0354 |  |  |
| TROUT | 0.687 | <1e-05 |  |  |
| USC MO | 0.229 | <1e-05 | 0.462 | <1e-05 |

**TABLE S5:**
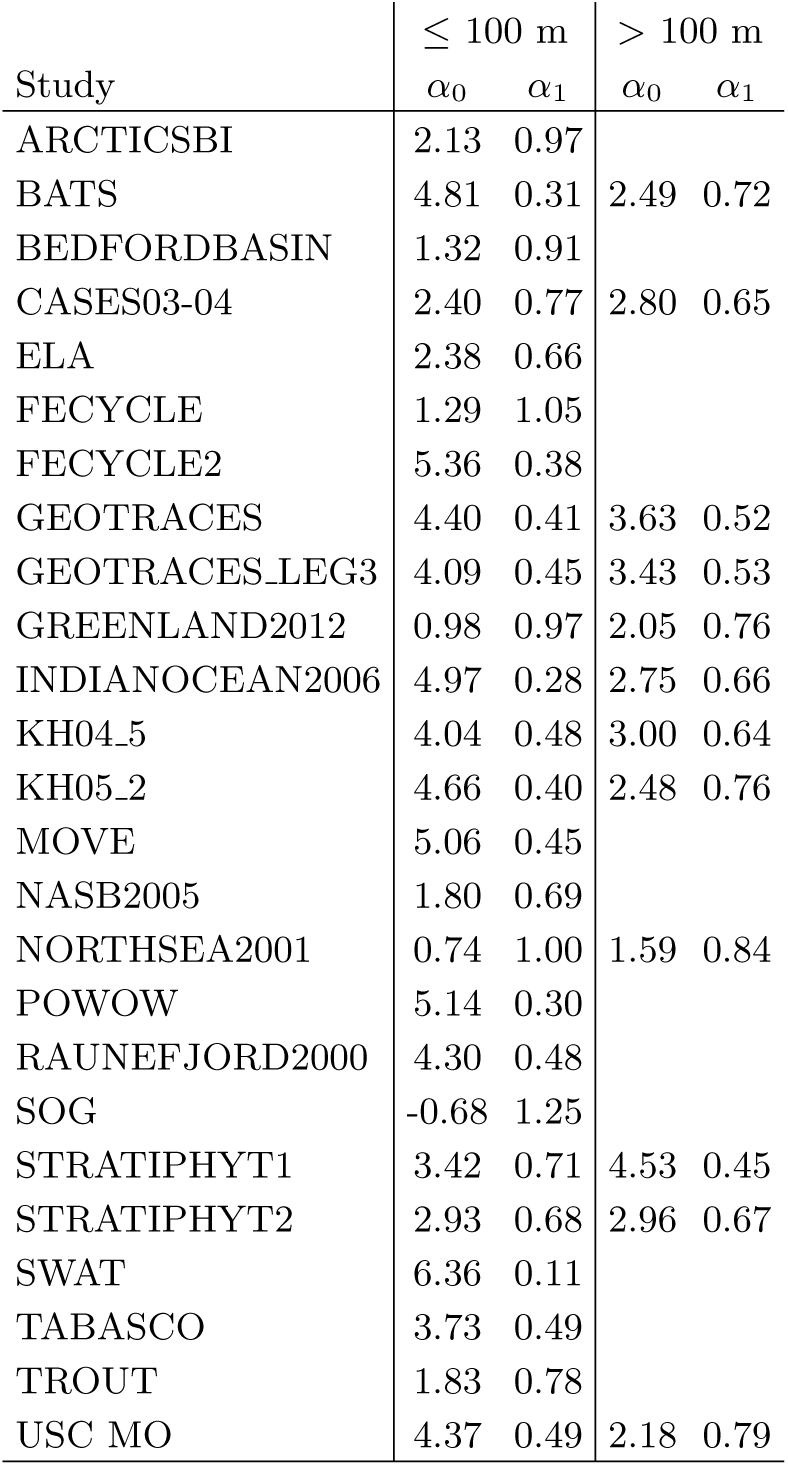
Power-law exponents, *α_1_*, and intercepts, *α_0_*, for each study from the mixed model allowing study-specific slopes and intercepts. Empty cells in a row denote the absence of samples collected at depths *>* 100 m for the study denoted in the left-most column.

| Study | $\leq 100$ m | | $> 100$ m | |
| --- | --- | --- | --- | --- |
| | $\alpha_0$ | $\alpha_1$ | $\alpha_0$ | $\alpha_1$ |
| ARCTICSBI | 2.13 | 0.97 |  |  |
| BATS | 4.81 | 0.31 | 2.49 | 0.72 |
| BEDFORDBASIN | 1.32 | 0.91 |  |  |
| CASES03-04 | 2.40 | 0.77 | 2.80 | 0.65 |
| ELA | 2.38 | 0.66 |  |  |
| FECYCLE | 1.29 | 1.05 |  |  |
| FECYCLE2 | 5.36 | 0.38 |  |  |
| GEOTRACES | 4.40 | 0.41 | 3.63 | 0.52 |
| GEOTRACES_LEG3 | 4.09 | 0.45 | 3.43 | 0.53 |
| GREENLAND2012 | 0.98 | 0.97 | 2.05 | 0.76 |
| INDIANOCEAN2006 | 4.97 | 0.28 | 2.75 | 0.66 |
| KH04.5 | 4.04 | 0.48 | 3.00 | 0.64 |
| KH05.2 | 4.66 | 0.40 | 2.48 | 0.76 |
| MOVE | 5.06 | 0.45 |  |  |
| NASB2005 | 1.80 | 0.69 |  |  |
| NORTHSEA2001 | 0.74 | 1.00 | 1.59 | 0.84 |
| POWOW | 5.14 | 0.30 |  |  |
| RAUNEFJORD2000 | 4.30 | 0.48 |  |  |
| SOG | -0.68 | 1.25 |  |  |
| STRATIPHYT1 | 3.42 | 0.71 | 4.53 | 0.45 |
| STRATIPHYT2 | 2.93 | 0.68 | 2.96 | 0.67 |
| SWAT | 6.36 | 0.11 |  |  |
| TABASCO | 3.73 | 0.49 |  |  |
| TROUT | 1.83 | 0.78 |  |  |
| USC MO | 4.37 | 0.49 | 2.18 | 0.79 |

**FIG. S1:**
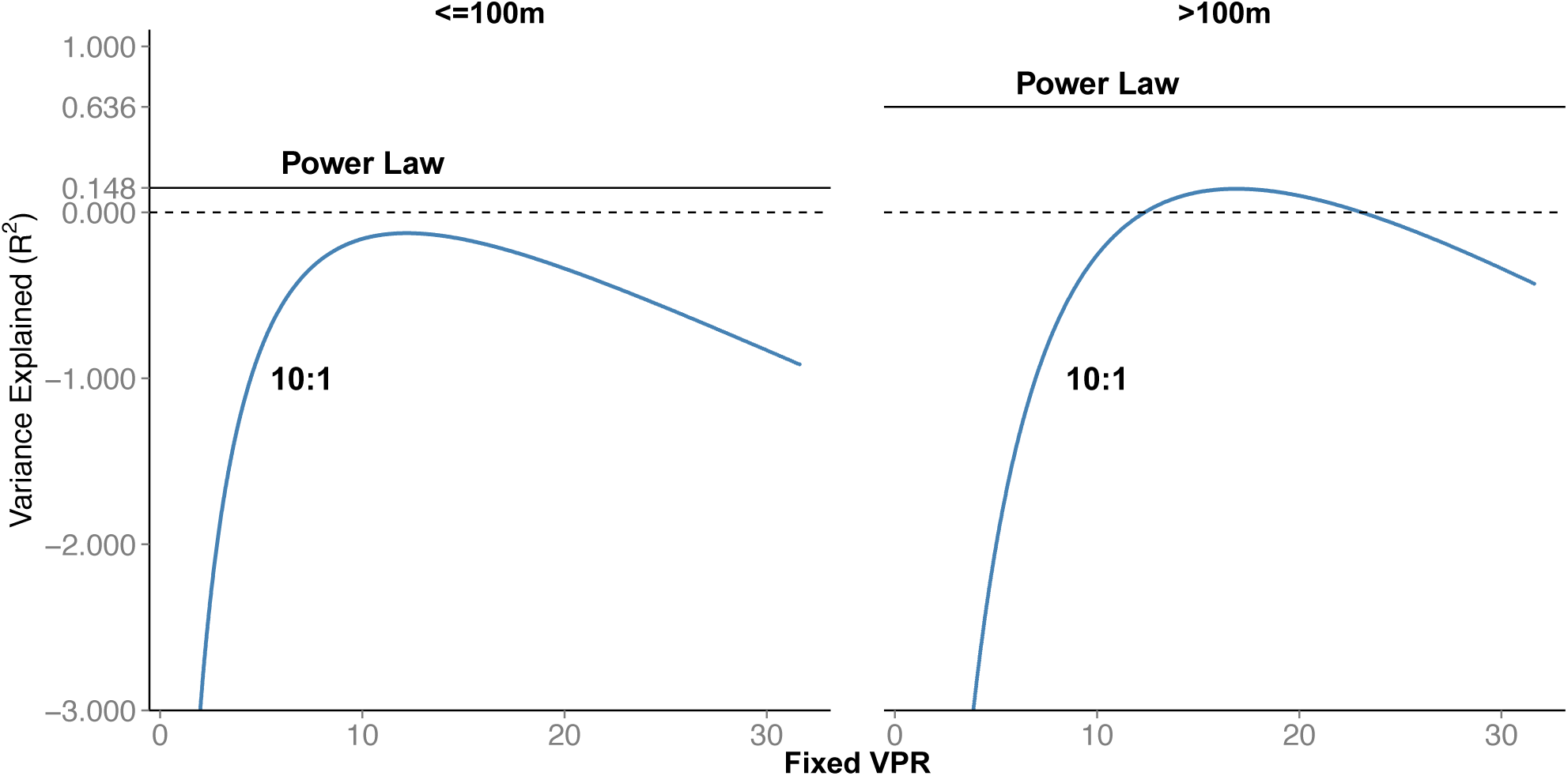
Explanatory power of fixed VPR models in the surface ocean (left) and deeper water column (right). The x-axis denotes the value *r* in the model *V* = *rP* where *V* denotes virus abundance and *P* denotes prokaryote abundance. The y-axis denotes the fraction of variance explained, *R*^2^. Here, *R*^2^ = 1 - SSE_model_*/*SSE_total_ where SSE_model_ is the sum of squared errors for the model and SSE_total_ is the sum of total squared errors.

**FIG. S2:**
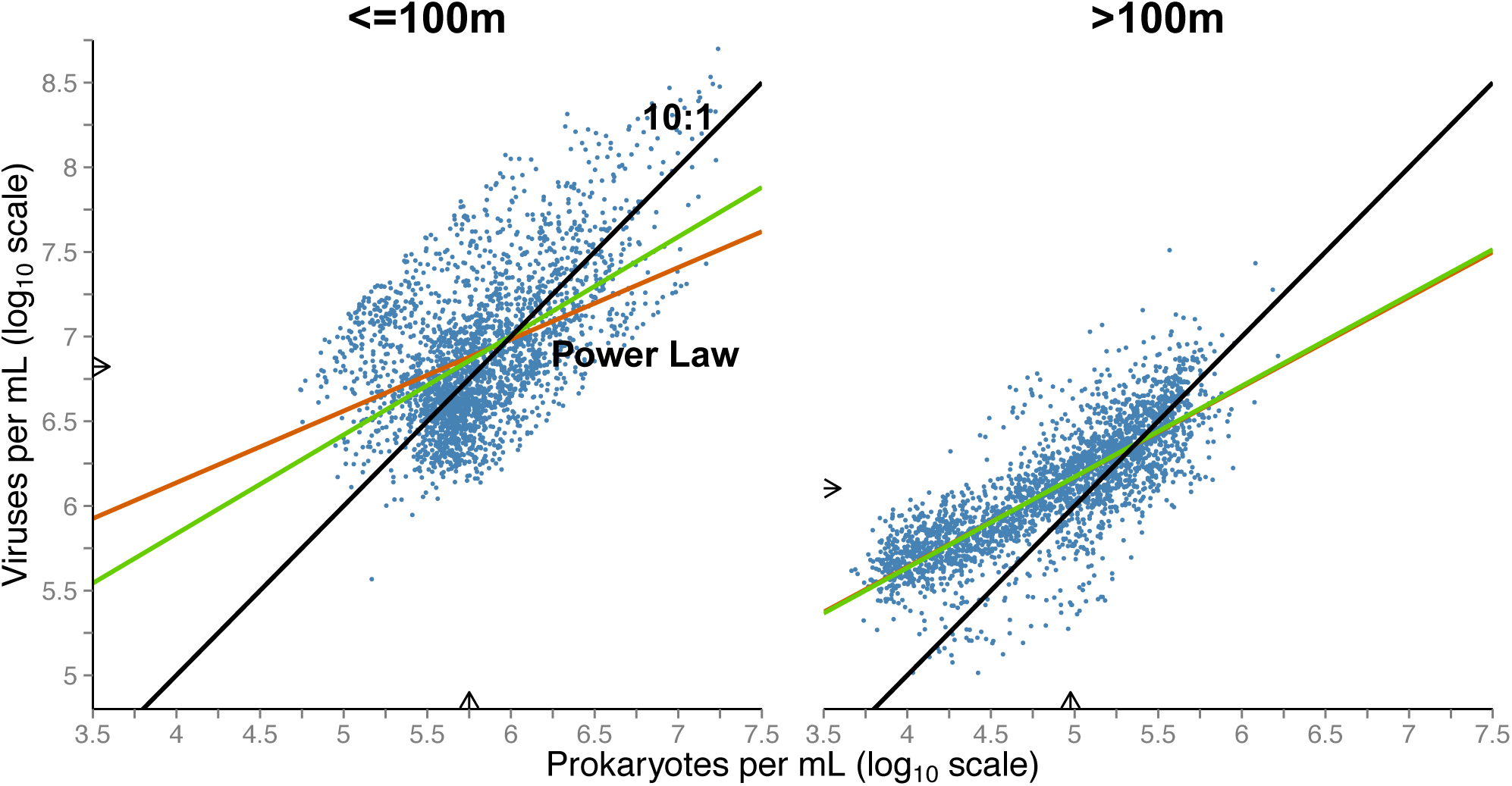
Explanatory power of fixed VPR models in the near-surface and sub-surface with and without outliers. The three lines in each panel denote the 10:1 line (black), power-law fit (red) and power-law fit when removing outliers (green). The *R*^2^ value for the power law fit for surface data excluding outliers is 0.30, has a slope of 0.58 and an intercept of 3.50. The *R*^2^ value for the power law fit for sub-surface data excluding outliers is 0.65, has a slope of 0.54 and an intercept of 3.49.

**FIG. S3:**
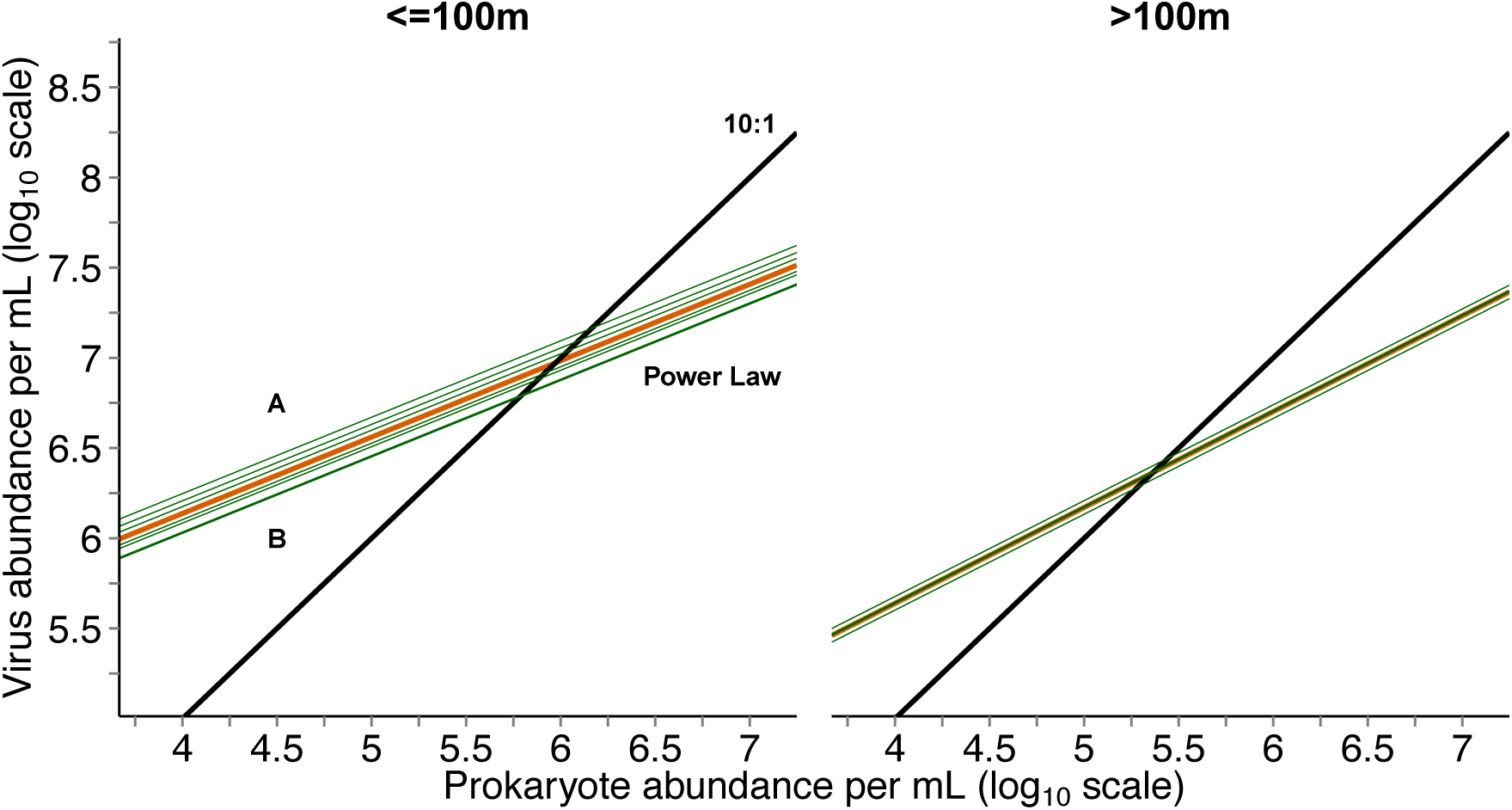
Constrained regression model for samples taken at depths *≤*100m (left) and *>* 100m (right) where the intercept for each study was permitted to vary (see Materials and Methods). Blue line denotes the 10:1 relationships, the red line denotes the best-fitting power-law model, and the remainder of lines denote the variable intercept model with intercept values reported in Table S3.

**FIG. S4:**
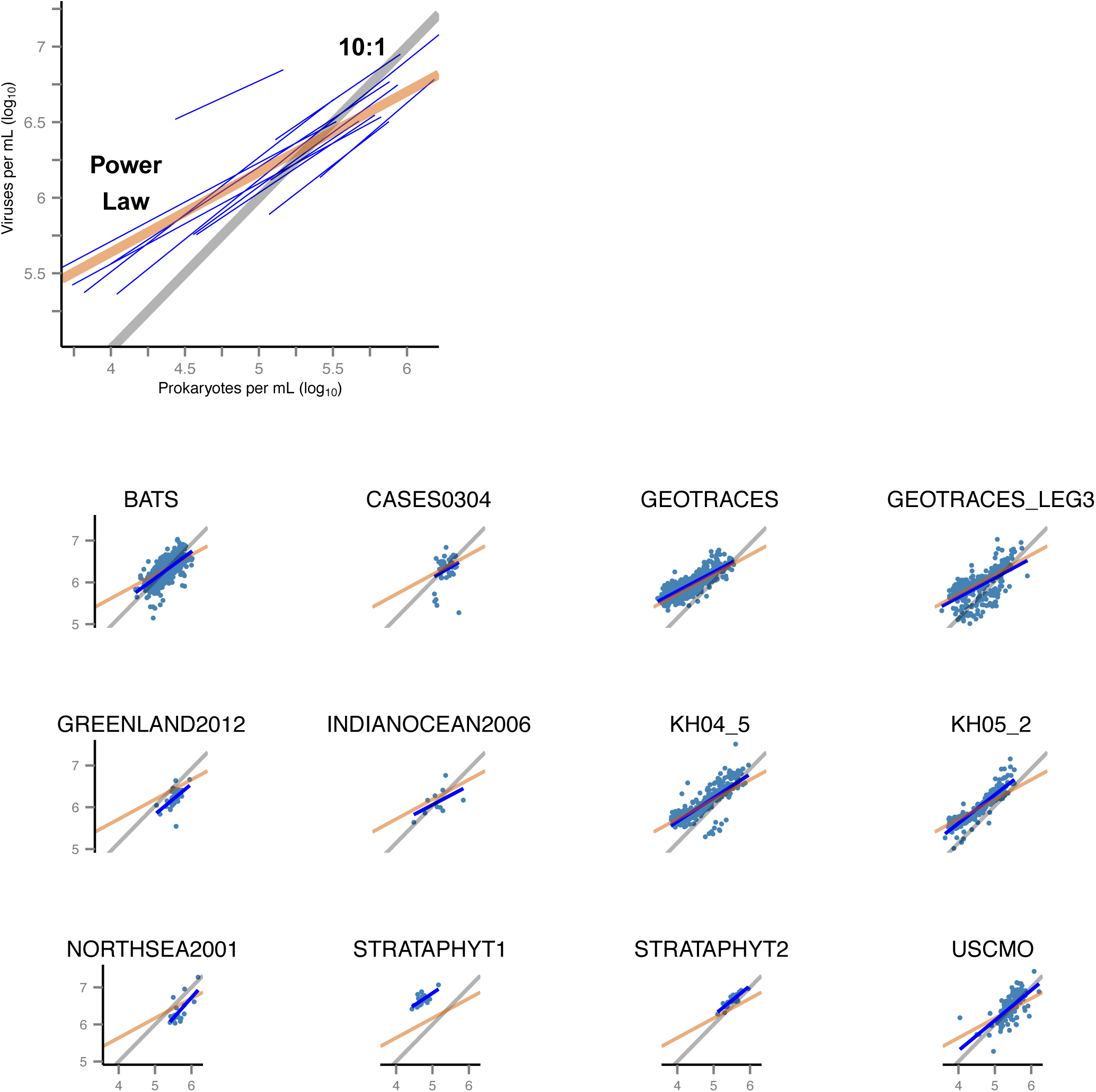
Virus-prokaryote relationships given the variable slope and intercept mixed-effects model for samples taken at depths greater than 100m. (Upper-left) Best-fit power-law for each study (blue lines) plotted along with the best-fit power-law of the entire dataset (red line) and the 10:1 line (grey line). (Individual panels) Best-fit power-law model (blue line) on log-transformed data (blue points) for each study, with the power-law model regression (red) and 10:1 line (black) as reference. The power-law exponents and associated confidence intervals are shown in Figure S5,

**FIG. S5:**
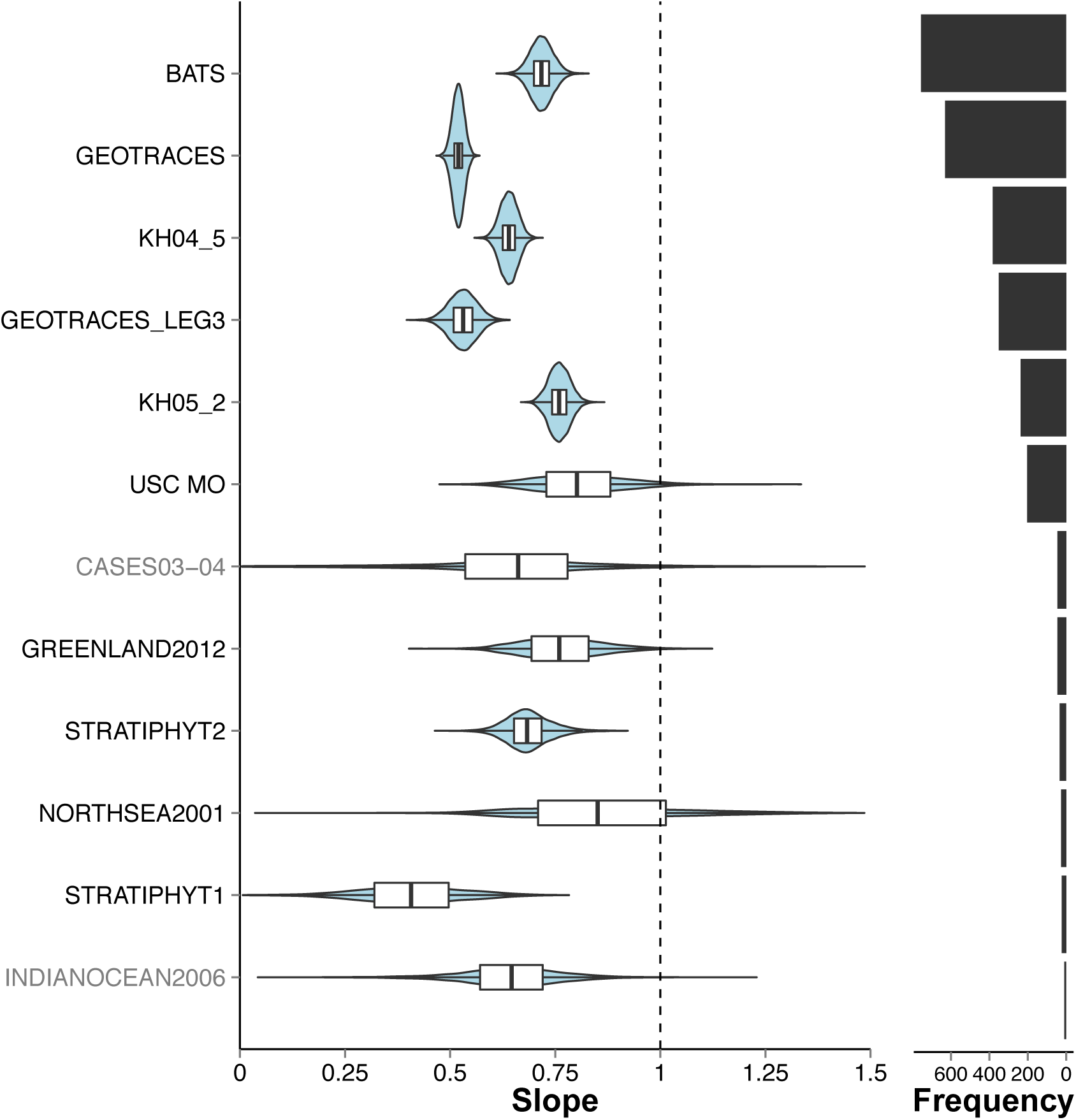
Study-specific 95% confidence intervals of power-law exponents for relationships between virus and prokaryote abundance from samples taken at depths greater than 100m. The confidence intervals are plotting using “violin” plots including the median (center black line), 75% distribution (white bars) and 95% distribution (black line), with the distribution overlaid (blue shaded area). The number of points included as part of each study is displayed on the right-most bar plots. Study labels in black indicate those studies whose linear regression had a p-value less than.05/12 while labels in gray indicate a p-value above this threshold.

